# Metazoan Orc6 Proteins Evolved Alternative Mechanisms for Association with the ORC Complex: Insights from *Drosophila* Modeling

**DOI:** 10.64898/2026.08.20.745992

**Authors:** Maxim Balasov, Etsuko Shibata, Katarina Akhmetova, Anindya Dutta, Igor Chesnokov

## Abstract

In eukaryotes, DNA replication requires the origin recognition complex (ORC), a six-subunit assembly that promotes replisome formation on chromosomal origins. Orc6 is the smallest and least evolutionarily conserved among all ORC subunits. In *Drosophila*, Orc6 binds tightly with the core ORC(1-5) and is required for DNA binding and replication initiation, whereas in *Xenopus* and human systems Orc6 loosely associates with the rest of the complex resulting in some differences for replication-associated activities. Despite these variations, Orc6 remains essential for viability in all species. In current study we analyzed specific residues within the C-terminal α11 helix that is critical for stable association of Orc6 with the ORC complex in *Drosophila.* Human Orc6 lacks these residues, however it possesses a strong nuclear localization signal (NLS) that is absent in *Drosophilidae.* We propose that this NLS drives human protein to the nucleus and compensates for weaker Orc6-ORC(1-5) interactions by increasing the nuclear concentration of Orc6 and shifting the equilibrium toward formation of the fully assembled ORC complex at the DNA.

## Introduction

The hexameric origin recognition complex (ORC) is an essential component of eukaryotic DNA replication. Originally discovered in *Saccharomyces cerevisiae*, subsequent studies in both yeast and higher eukaryotes established the foundation for understanding the functions of this critical initiation factor functions (BELL AND STILLMAN 1992), (BELL 2002), (SASAKI AND GILBERT 2007), (CHESNOKOV 2007), (DUNCKER *et al*. 2009). ORC binds replication origin sites in an ATP-dependent manner and directs assembly of the pre-replicative complex (pre-RC) at these origins (BELL 2002) (MACHIDA *et al*. 2005a). ORC complexes have since been identified in numerous metazoan species (BELL 2002), (CHESNOKOV 2007). Among all ORC subunits, Orc6 is the least evolutionary conserved and most enigmatic. In *S. cerevisiae*, Orc6 is not essential for DNA binding but is required for cell viability (LI AND HERSKOWITZ 1993), (NEWLON 1993), (LEE AND BELL 1997) and maintenance of the pre-RC, specifically through recruitment of Cdt1 followed by MCM loading (SEMPLE *et al*. 2006), (CHEN *et al*. 2007). In contrast, *Schizosaccharomyces pombe* and metazoan Orc6 proteins (CHESNOKOV *et al*. 1999), (MOON *et al*. 1999), (DHAR AND DUTTA 2000) are similar in size and are considerably smaller than *S. cerevisiae* Orc6. In *Drosophila melanogaster*, Orc6 is essential for ORC-dependent DNA binding and DNA replication (CHESNOKOV *et al*. 2001), (BALASOV *et al*. 2007). In *Xenopus* and human systems, Orc6 loosely associates with the ORC1–5 core complex, and some studies suggest that Orc6 activities in different species are divers (GILLESPIE *et al*. 2001), (VASHEE *et al*. 2001) (VASHEE *et al*. 2003). This apparent discrepancy may reflect differences in the affinity of Orc6 for the ORC1–5 core complex among distant metazoan species. Nevertheless, in both *Drosophila* and humans, Orc6 is essential for survival, and mutations in Orc6 result in severe developmental defects leading to Meier-Gorlin syndrome and lethality in most severe cases (BALASOV *et al*. 2009), (BALASOV *et al*. 2015), (SHALEV *et al*. 2015), (BALASOV *et al*. 2020). Structural studies demonstrated that Orc6 interacts with the rest of the ORC through a C-terminal α-helix (BLEICHERT *et al*. 2013), (BLEICHERT *et al*. 2015), (XU *et al*. 2020). This helical structure contains conserved amino acid residues that are identical in human and *Drosophila* Orc6 proteins (BALASOV *et al*. 2009), (BALASOV *et al*. 2015), (BLEICHERT *et al*. 2013) and mediate interaction with the complex through the Orc3 subunit (BLEICHERT *et al*. 2013), (BLEICHERT *et al*. 2015).

In the current study, we used humanized *Drosophila* models to decipher molecular mechanisms defining diverse Orc6 behavior in flies and humans. Specifically, we investigated the function of the *Drosophila* Orc6 C-terminus *in vivo* and found that the Methionine residue at position 232 is important for *Drosophila* Orc6 tight association with the ORC complex. Notably, human ORC6 lacks this Methionine residue. However, in humans ORC6 possesses a strong nuclear localization signal (XU *et al*. 2020). We hypothesize that a strong NLS found in human Orc6 increases the nuclear concentration of the protein, thereby shifting the equilibrium toward ORC complex assembly at the origin and compensating for the lower affinity of the human Orc6 subunit for the core complex.

## Results

Structural studies of metazoan ORC revealed that Orc6 protein integrates into the complex through interaction with the Orc3 subunit via its α11 helix (BLEICHERT *et al*. 2013), (BLEICHERT *et al*. 2015), (XU *et al*. 2020). The N-terminal region of this helix is highly conserved between human and *Drosophila* (**Figure 1**). Mutations in this region, highlighted in **Figure 1**, disrupt Orc6 association with the complex and are known to contribute to disease development (BALASOV *et al*. 2009), (BLEICHERT *et al*. 2013), (BALASOV *et al*. 2015). In contrast, the remaining portion of the α11 helix is less conserved and may play an important role in strengthening Orc6 association with the complex. To further investigate the differences between metazoan Orc6 C-terminal domains that mediate stable association with the Orc1–5 complex, we performed Basic Local Alignment Search Tool (BLAST) analysis across different *Drosophila* species. Our analysis also revealed that *Drosophila melanogaster* contains a unique C-terminal tail extension QSHMDSQLLEA that is absent in mammals (**Figure 1**). In this study, we generated a series of human–*Drosophila* Orc6 hybrids including point mutants to identify the amino acid determinants within the *Drosophila* Orc6 C-terminus required for stable association with the ORC complex *in vivo*. We employed three complementary approaches to evaluate the effects of these mutations: (1) interaction of Orc6 with the ORC complex measured by co-immunoprecipitation; (2) the ability of mutant proteins to rescue the *orc6* deletion phenotype, and (3) analysis of mitotic chromosome integrity as an indicator of successful DNA replication.

**Figure 1.**
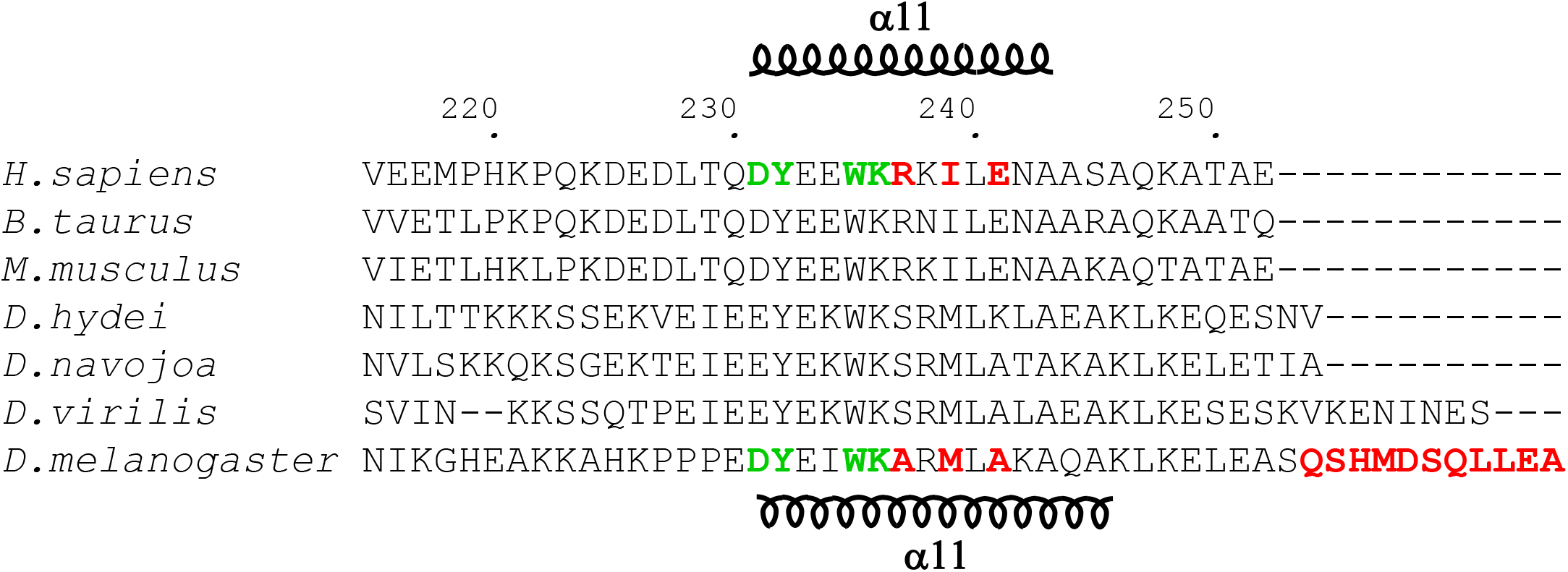
Amino acid sequence alignment. The C-terminal tails of the Orc6 proteins from three species of mammals are aligned with four species of *Drosophila.* Amino acids studied in previous work are shown in green (Balasov et al., 2009, 2015; Bleichert et al., 2013). Amino acids investigated in the current study and *D.melanogaster* C-terminus extension are shown in red. Predicted a11 helix in *Drosophila* and human proteins are shown in coils.

First, we created the human-*Drosophila* Orc6 hybrid (Orc6Hs(243)Dm(22)), which contains the first 243 amino acids of the human protein and 22 amino acids of the *Drosophila* protein (**Figure 2**). This construct includes human α11 helix together with the *Drosophila* C-terminal tail extension QSHMDSQLLEA. Functional analysis revealed that, similar to the human transgene, this chimeric protein did not associate with the ORC complex (**Figures 2** and **4A**), did not rescue the *orc6* deletion phenotype (**Figure 2**) and exhibited severe chromosome defects that resembled the effect of *orc6* deletion (**Figures 5A, B**). These findings suggest that the *Drosophila* Orc6 C-terminal extension (QSHMDSQLLEA) is dispensable for ORC complex association. Therefore, removal of this extension from Drosophila Orc6 is unlikely to have a significant impact on Orc6 activity or its interaction with the ORC complex. To confirm this conclusion, we deleted the C-terminal extension QSHMDSQLLEA from *Drosophila* Orc6 and found that the Orc6Dm(246) deletion mutant was functionally identical to the wild-type *Drosophila* Orc6 in all assays tested, including ORC complex association (**Figure 4B**), rescue of the Orc6 deletion phenotype (**Figure 2**), and chromosome integrity (**Figures 5A,B**). Next, we generated the Orc6Hs(236)Dm(28) construct, which contains 236 amino acids from the human protein and C-terminal 28 amino acids from the *Drosophila* protein (**Figure 2**). Specifically, this chimeric construct contains the conserved N-terminal residues of the human α11 helix (DYEEWK) fused to the variable C-terminal residues of the *Drosophila* α11 helix (ARMLAKAQA) (**Figure 2**). In effect, the Orc6Hs(236)Dm(28) combines the *Drosophila* α11 helix together with the C-terminal tail extension (QSHMDSQLLEA). This hybrid Orc6 associated with the ORC complex (**Figure 4B**), rescued the lethality caused by Orc6 deletion (**Figure 2**), and restored mitotic chromosome integrity (**Figure 5A, B**). Given that the C-terminal tail extension is not required for ORC complex association, we deleted this region to generate Orc6Hs(236)Dm(17). Functional analysis revealed that Orc6Hs(236)Dm(17) retained activity comparable to that of Orc6Hs(236)Dm(28) (**Figures 2**, **4C, and 5B**). Taken together, these findings identify the *Drosophila* α11 helix as the critical determinant mediating stable association of the chimeric Orc6 protein with the ORC complex, while demonstrating that the C-terminal tail extension, specific for *D. melanogaster*, does not make a significant contribution to this interaction.

**Figure 2.**
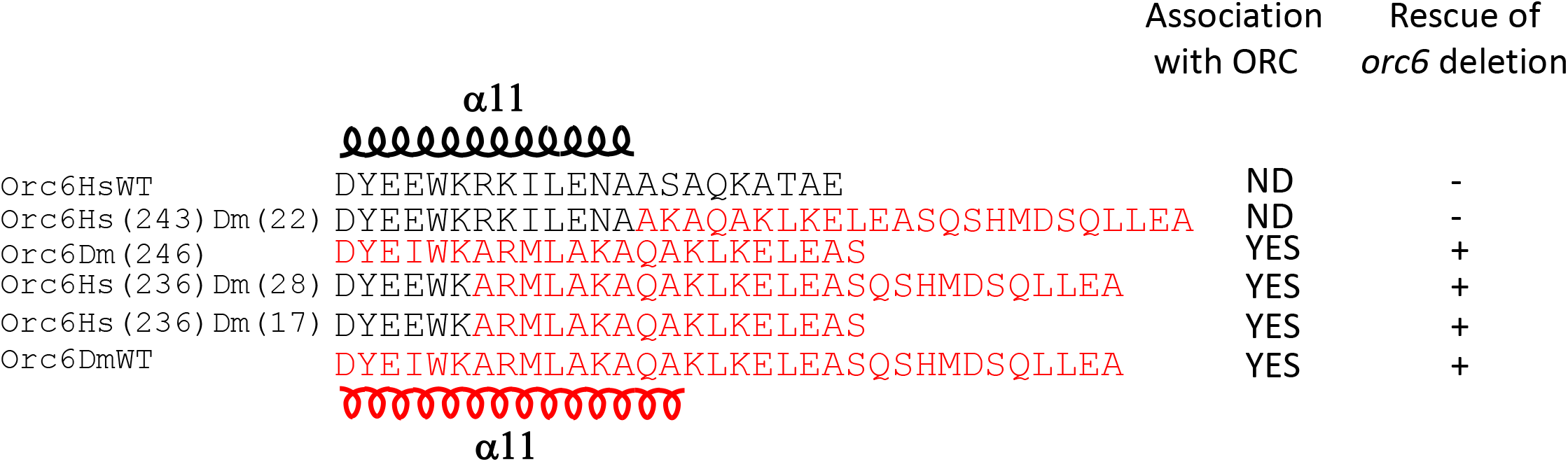
Effect of the α11 helix and tail extension on Orc6 association with *Drosophila* ORC complex. In the hybrid proteins Orc6Hs(243)Dm(22), Orc6Hs(236)Dm(28), and Orc6Hs(236)Dm(17), the human-derived regions are shown in black and the *Drosophila*- derived regions in red. The numbers in parentheses indicate the number of amino acids contributed by the human or *Drosophila*. For example, Orc6Hs(243)Dm(22) indicates that the first 243 amino acids are from the human sequence, while the remaining 22 amino acids are from *Drosophila*. Orc6Dm(246) - *Drosophila* Orc6 missing C-terminal extension. The predicted α11 helix in both human and *Drosophila* is depicted as coils. ND - not detected. ‘+’ rescue, ‘-’ – no rescue.

**Figure 3.**
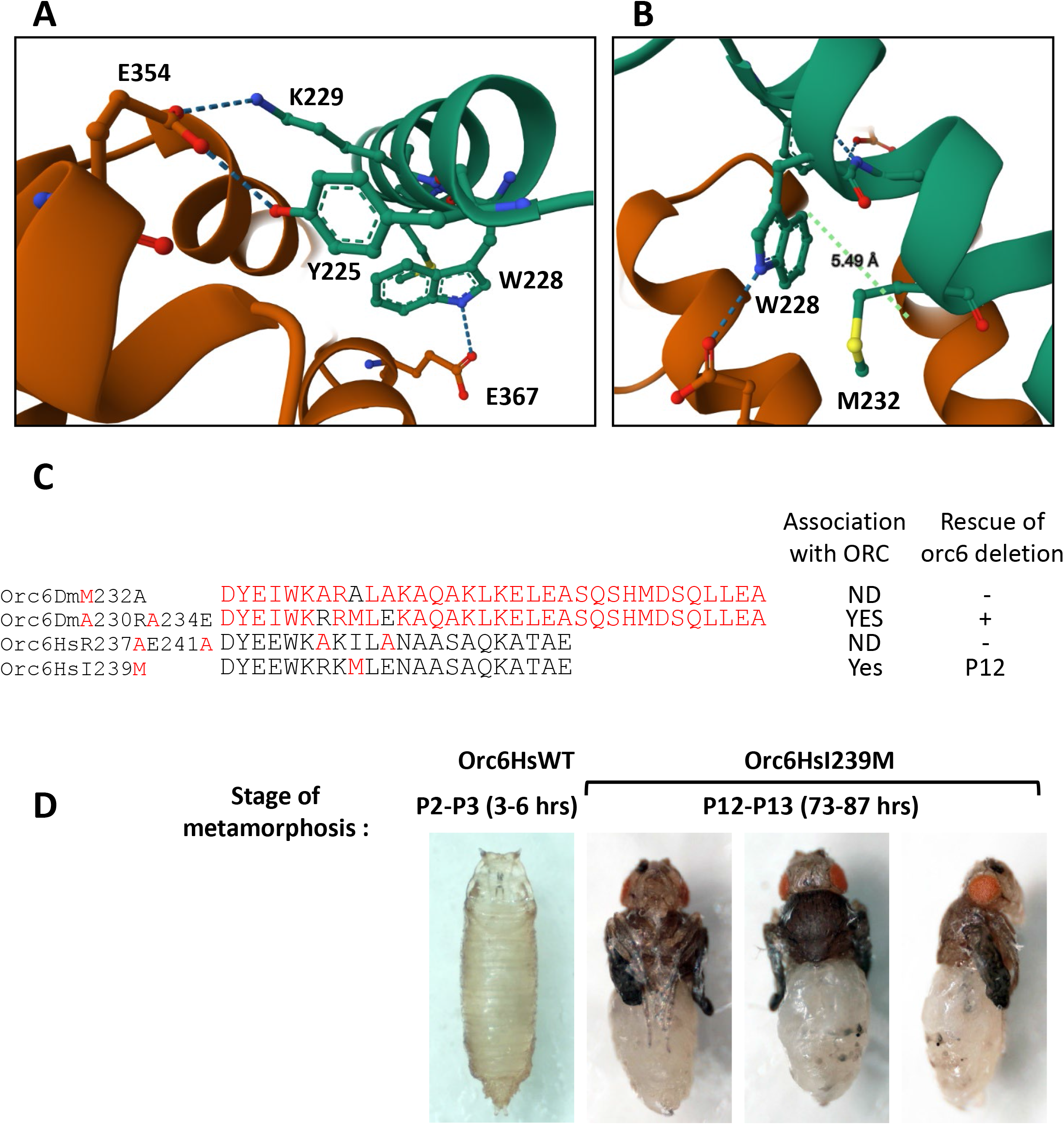
Effects of amino acid mutations in the α11 helix on Orc6 association with the *Drosophila* ORC complex. (A,B) AlphaFold-predicted hydrogen bonds between *Drosophila* Orc6 and Orc3. The Orc6 α11 helix is shown in green, and Orc3 is shown in brown. **(C)** Alignment of the C-terminus of human and *Drosophila* Orc6. Human amino acids are shown in black, and *Drosophila* sequences are shown in red. ND - not detected. “+” indicates rescue to adulthood, “-” indicates no rescue, P12 - rescue to pupal stage 12. **(D)** Orc6HsWT – rescued with human wild-type Orc6. Orc6HsI239M – human Orc6 in which Isoleucine at position 239 is replaced by Methionine.

**Figure 4.**
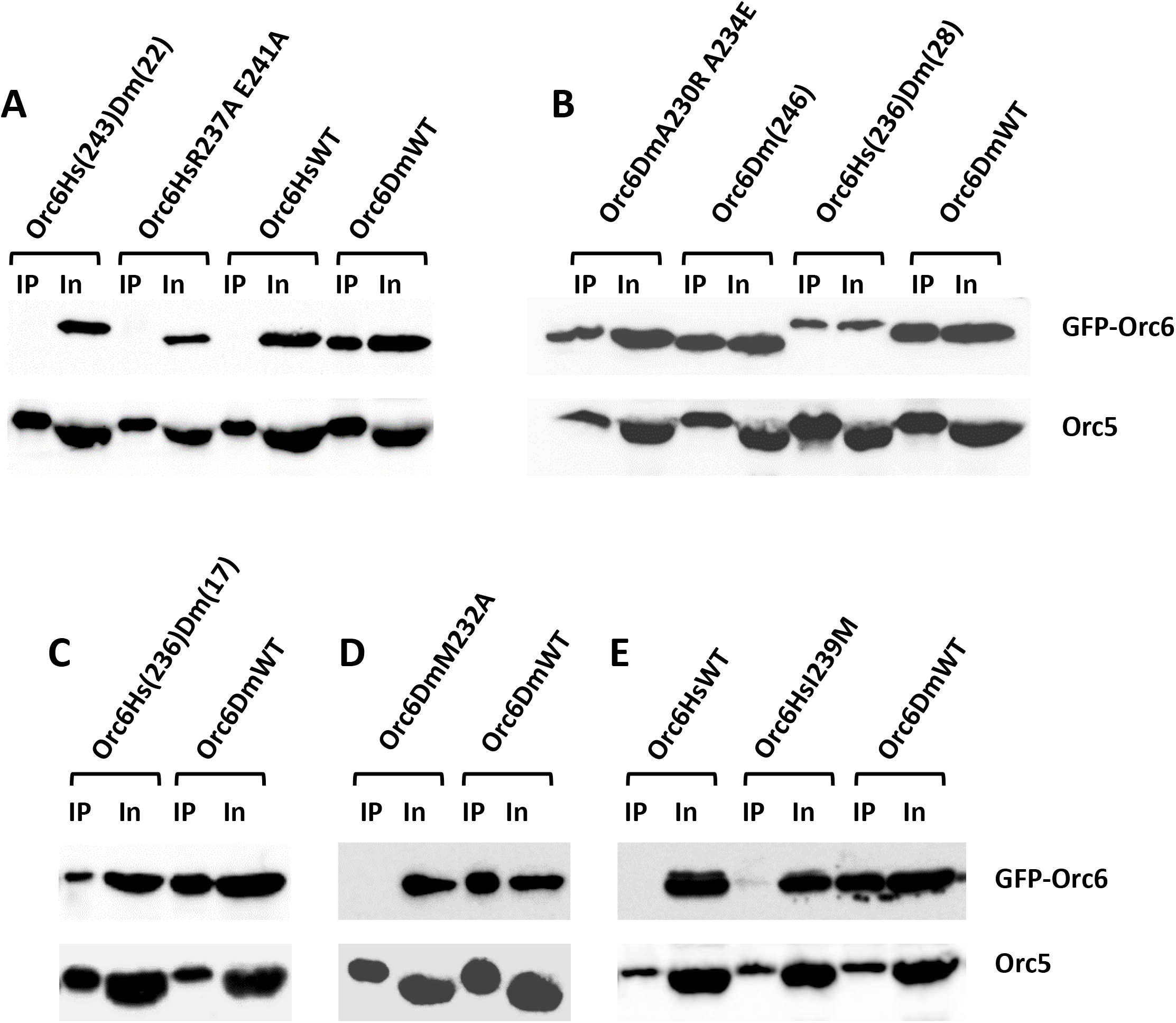
Association with the ORC complex. (A,B,C,D,E) The ORC complex was immunoprecipitated from ovaries expressing GFP-tagged Orc6 transgenes using anti-Orc2 antibodies. In the hybrid proteins Orc6Hs(243)Dm(22), Orc6Hs(236)Dm(28), and Orc6Hs(236)Dm(17), the numbers in parentheses indicate the number of amino acids contributed by the human or *Drosophila* sequences. Orc6HsWT refers to human wild-type Orc6 (negative control), and Orc6DmWT refers to *Drosophila* wild-type Orc6 (positive control). Point mutations are indicated using single-letter abbreviations; for example, Orc6HsI239M indicates that isoleucine at position 239 in human Orc6 was replaced by methionine.

**Figure 5.**
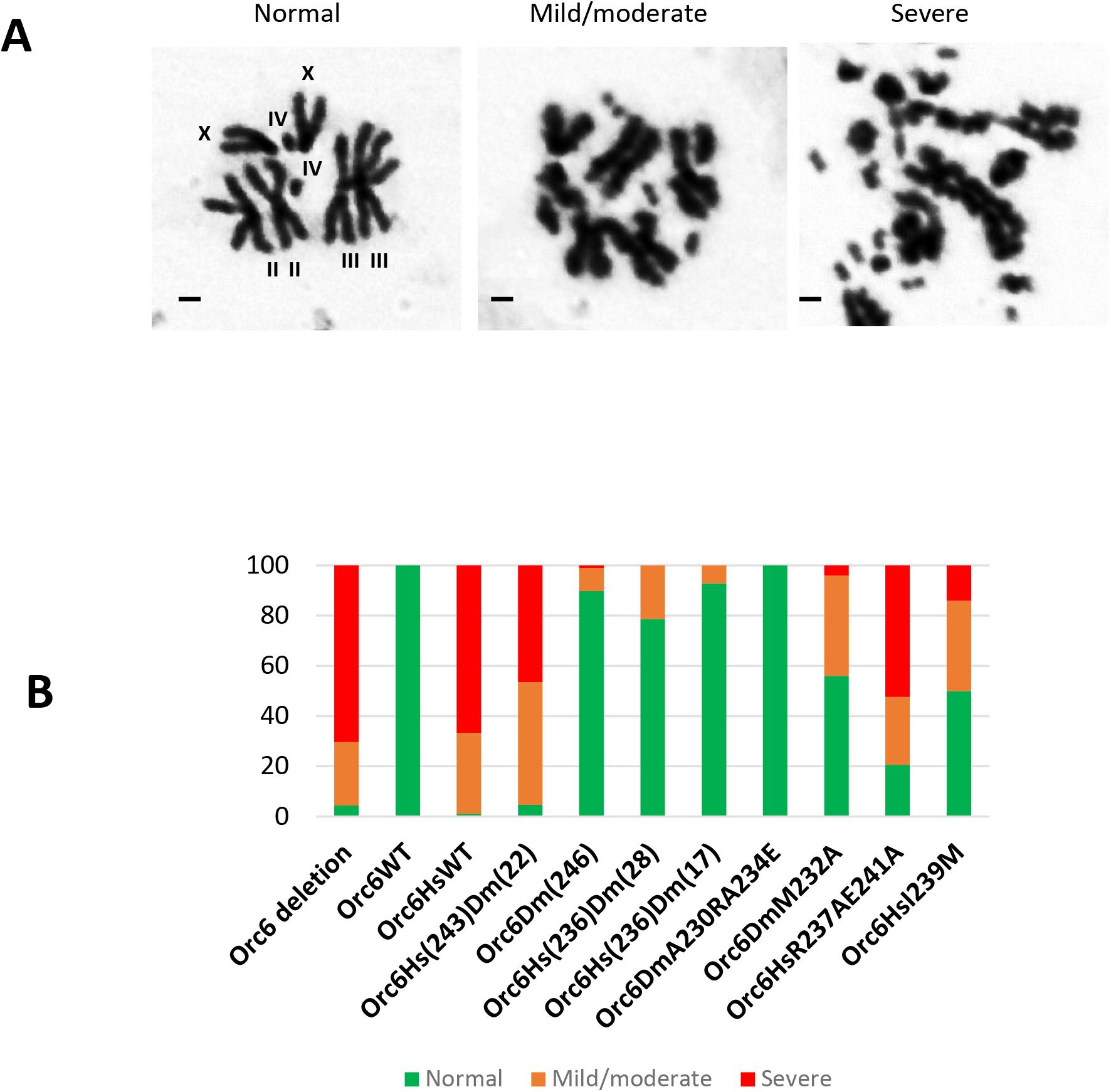
Mitoses in larval neuroblasts. **(A)** The karyotypes were divided onto three classes. Normal – all chromosomes are clearly visible, no defects. Mild/moderate - karyotypes are defective, some chromosomes can be identified, but contain rearrangements (deletions, duplications, bridges). Severe - chromosomes are abnormally condensed and fragmented. “XX” - pair of the X, “II II” – second, “III III”- third, and “IV IV” - fourth chromosomes. Scale bar 1µm. **(B)** Percentage of normal (green) and defective (orange and red) karyotypes was plotted on a bar chart. Orc6DmWT represent positive control. Orc6 deletion and Orc6HsWT represent negative control.

The N-terminal part of the α11 helix is nearly identical in human and *Drosophila* Orc6 proteins; however, in humans Orc6 loosely associates with the ORC complex (Vashee *et al*. 2001), (Vashee *et al*. 2003), (Ranjan and Gossen 2006). This suggests that the C-terminal region of the *Drosophila* α11 helix contains specific features that stabilize interaction with the complex. To investigate this further, we modeled the Orc3-Orc6 interaction using AlphaFold (https://www.cosmic2.org/alphafold2). In addition to the well-characterized Y225 and W228 residues which form hydrogen bonds with Orc3 (**Figure 3A**), we identified M232 positioned at a distance favorable for an S-aromatic interaction with W228 (**Figure 3B**) (ALEDO 2019), (VALLEY *et al*. 2012). To test the importance of M232, we generated the Orc6DmM232A mutant and found that it failed to associate with the ORC complex in *Drosophila* (**Figure 4D**), was unable to rescue the *orc6* deletion phenotype in fly (**Figure 3C**), and the mitotic chromosome integrity in cells carrying this mutant was significantly reduced (**Figure 5A, B**). M232 is flanked by the non-conserved residues A230 and A234, which correspond to R237 and E241 in human Orc6, respectively (**Figure 1**, **Figure 3C**). To assess whether these substitutions contribute to species-specific differences, we replaced A230 with R and A234 with E, generating the Orc6DmA230R/A234E mutant. However, this mutant performed indistinguishably from wild type DmOrc6WT in all tested assays (**Figures 3C**, **4B, and 5B**). Conversely, to determine whether introduction of the *Drosophila*-specific residues A230 and A234 could enhance the function of human Orc6 in *Drosophila*, we generated the Orc6HsR237A/E241A mutant and assessed its activity. Consistent with human Orc6WT, the Orc6HsR237A/E241A mutant failed to associate with the ORC complex (**Figures 4A**), did not complement the *orc6* deletion phenotype (**Figure 3C**), and was unable to restore chromosome integrity (**Figure 5B**). In human ORC6, I239 corresponds to *Drosophila* M232 and is positioned at a similar distance from the conserved W235 within the α11 helix as M232 is from W228 in flies (**Figure 1**). Because M232 contributes to Orc6 function in *Drosophila*, we tested whether an I239 to M239 substitution could improve human Orc6 activity in *Drosophila* by generating and analyzing the Orc6HsI239M mutant. Remarkably, this single substitution supported fly development through metamorphosis up to stages P12– P13, which corresponds to nearly fully developed adult flies (**Figure 3D**). This phenotype correlated with detectable ORC complex association (**Figure 4E**) and substantial restoration of mitotic chromosome integrity (**Figure 5B**). By contrast, wild-type human Orc6HsWT does not associate with the ORC complex (**Figure 4A, E**), fails to rescue the *orc6* deletion phenotype, resulting in lethality at the same developmental stage as *orc6* deletion mutants (**Figure 3D**) and does not restore mitotic chromosome integrity (**Figure 5B**). Collectively, the beneficial effect of the Methionine substitution in human Orc6 suggests that *Drosophila* M232 is a critical determinant that promotes stable Orc6 association with the ORC complex in flies.

Despite millions of years of separate evolution, distantly related *Drosophila* species retain Methionine at this conserved position, whereas mammals carry Isoleucine instead (**Figure 1**). Therefore, we investigated the effect of Orc6HsI239M mutation in human cells. We transfected HCT116 cells with the Orc6HsI239M transgene and compared its activity with Orc6HsWT (wild type, positive control) and the previously characterized Meier-Gorlin syndrome mutant Orc6HsY232S, whose ability to bind the ORC complex is significantly reduced, and therefore it served as a negative control (BLEICHERT *et al*. 2013), (BALASOV *et al*. 2015). Pull-down experiments using cell extracts showed that the Orc6HsI239M mutant did not exhibit enhanced association with the ORC complex relative to wild-type Orc6. However, its binding to the complex was much stronger than that of the Orc6HsY232S MGS mutant (**Figure 6A**). To test whether Orc6HsI239M could functionally substitute for Orc6HsWT, we performed an MTT 3-(4, 5- dimethylthiazolyl-2)-2, 5-diphenyltetrazolium bromide) assay to assess cell proliferation. We found that the Orc6HsI239M mutant successfully restored cell proliferation to a level comparable to that of Orc6HsWT (**Figure 6B**), indicating that the I239M substitution is not detrimental to the essential cellular functions of human Orc6.

**Figure 6.**
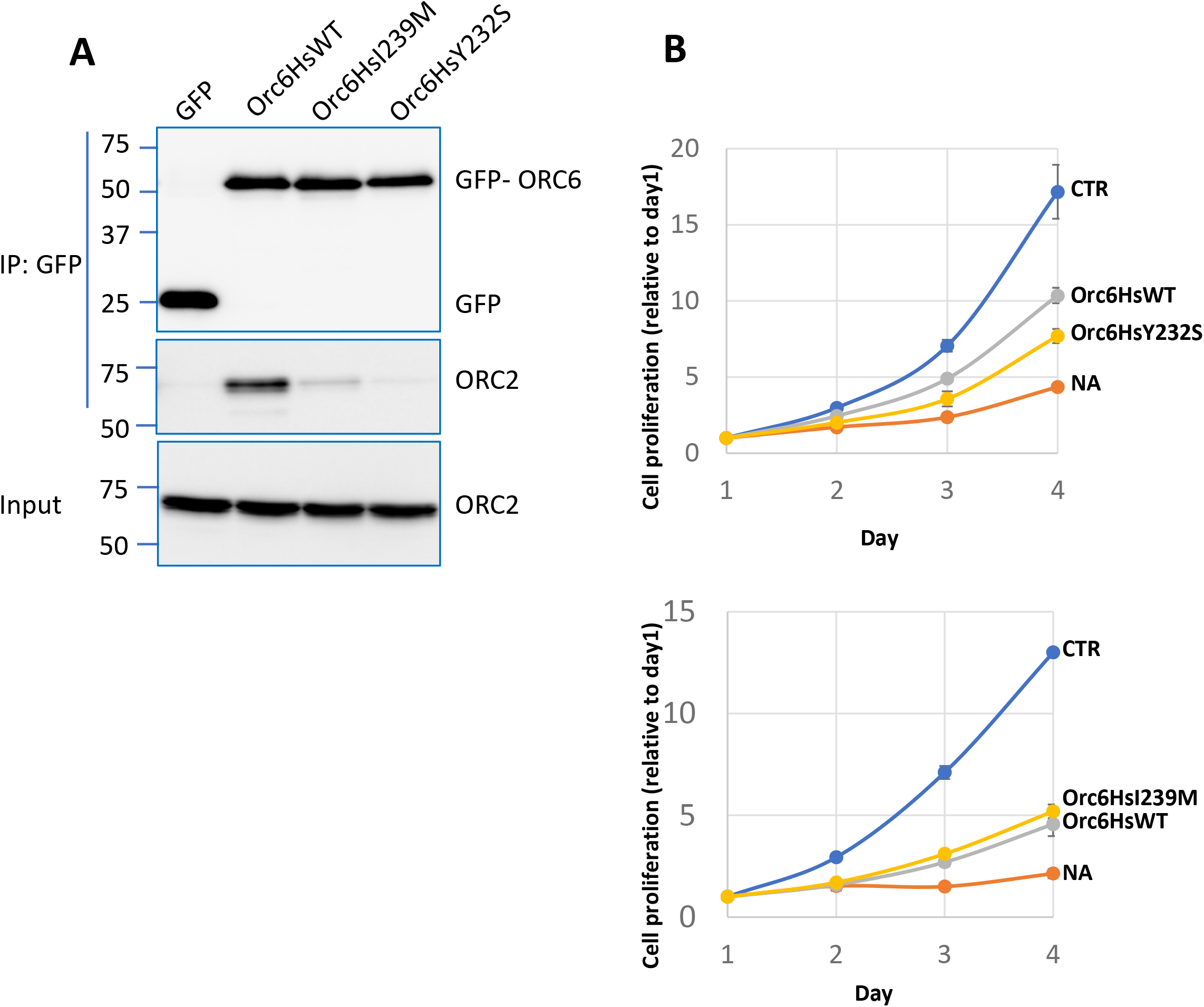
Interaction of human Orc6 with the ORC complex and cell proliferation in HTC116 cells. **(A)** Co-immunoprecipitation of the wild type human Orc6 (Orc6HsWT), methionine mutant (Orc6HsI239M) and Meier–Gorlin mutant (Orc6HsY232S) with core subunit Orc2. Orc6HsI239M indicates that Isoleucine at position 239 in human Orc6 was replaced by Methionine. Orc6HsY232S – Tyrosine at position 232 replaced with Serine. **(B)** MTT assay evaluating cell proliferation over 4 days in cells expressing human WT or mutant ORC6 following depletion of endogenous ORC6. CTR represents control cells, and NA indicates knockdown condition without expression of GFP-HsORC6.

In our previous study, we showed that elevated expression of human Orc6 in *Drosophila* promotes its association with the *Drosophila* ORC complex, whereas expression under the control of the native *orc6* promoter does not (BALASOV *et al*. 2020). These findings suggest that an increased local concentration of Orc6 can shift the equilibrium toward formation of the fully assembled complex. We propose that the presence of a strong nuclear localization signal may enhance the association of Orc6 with the human ORC(1–5) complex by increasing the intranuclear concentration of Orc6. Indeed, in our structural studies of human ORC6 (XU *et al*. 2020) we identified amino acid residues R198/K199/R200/K201 which are important for DNA binding. The same amino acids are also predicted to constitute a strong nuclear localization signal (NLS). In the current work, we used the web-based NLS Mapper tool (KOSUGI *et al*. 2009b) to predict NLS activity for the species shown in **Figure 1**. ORC6 proteins from *H. sapiens*, *B. taurus*, and *M. musculus* possess a strong nuclear localization signal, however species of *Drosophila* family do not (**Supplementary Figure 1, 2**). Although NLS Mapper is considered highly accurate in predicting nuclear localization activity (KOSUGI *et al*. 2009a), we experimentally validated these predictions by expressing Orc6HsWT, Orc6DmWT, and the NLS mutant Orc6HsR198A/K199A/R200A/K201A proteins in both *Drosophila* and human cells. Consistent with the prediction, human ORC6 (Orc6HsWT) is located exclusively to the nucleus in both *Drosophila* and human cells (**Figure 7**). In contrast, *Drosophila* Orc6 (Orc6DmWT) was detected in both the cytoplasm and nucleus (**Figure 7**) consistent with our previous analyses (CHESNOKOV *et al*. 2001), (CHESNOKOV *et al*. 2003), (HUIJBREGTS *et al*. 2009), (AKHMETOVA *et al*. 2015).

**Figure 7.**
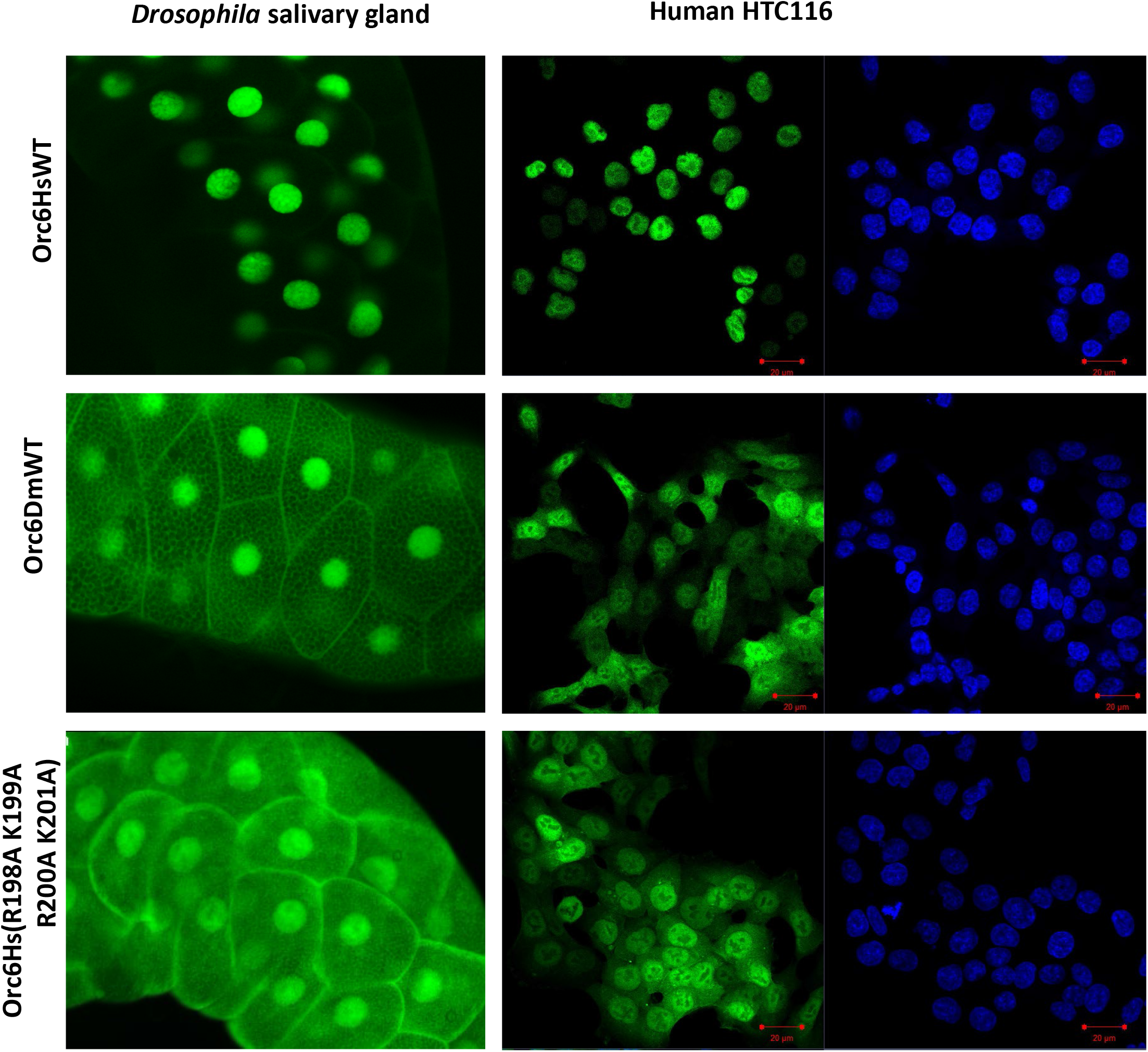
Nuclear localization of human and *Drosophila* Orc6. GFP-tagged Orc6 transgenes were expressed in *Drosophila* salivary glands or in human HTC116 cells. Orc6HsWT - wild type human Orc6, Orc6DmWT - wild type *Drosophila* Orc6, Orc6Hs(R198A,K199A,R200A,K201A) - human Orc6 with mutated nuclear localization signal.

Mutation of the NLS in human Orc6 (HsR198A/K199A/R200A/K201A) resulted in redistribution of Orc6 between the nucleus and cytoplasm (**Figure 7**). These results suggest that a strong NLS increases the nuclear concentration of Orc6, thereby promoting conditions favorable for its association with the DNA and the assembly of the functional ORC complex at the origin.

## Discussion

Even though Orc6 is an essential component for the initiation of DNA replication, its functions vary in different eukaryotic species. In yeast *S. cerevisiae*, Orc6 is an integral part of the ORC complex (LI *et al*. 2018). *Drosophila* Orc6 is not homologous in sequence to the budding yeast protein, but similarly to the budding yeast Orc6 it is tightly associated with other ORC subunits. In *Drosophila*, Orc6 binds DNA and may participate in positioning of ORC at the origin. Human Orc6 is homologous with *Drosophila* protein, it also possesses DNA binding ability and required for replication (BALASOV *et al*. 2009), (LIU *et al*. 2011), (PRASANTH *et al*. 2002), (THOMAE *et al*. 2008), (XU *et al*. 2020). However, human Orc6 loosely associates with core ORC subunits. The severe disruption of Orc6 results in lethality in both humans and *Drosophila* (BALASOV *et al*. 2009), (SHALEV *et al*. 2015). This indicates that the ORC complex critically depends on Orc6 for proper functioning.

According to the law of mass action, the rate of complex assembly is directly proportional to both the affinity between subunits and their local concentration. These two distinct mechanisms shift the equilibrium toward complex formation through different physical principles. *Drosophilidae* appear to have evolved stronger intermolecular interactions in part by acquiring M232 in addition to the conserved Y225 and W228 residues in alpha11 helix. In our previous studies, we tested these residues in a live *Drosophila* model and demonstrated that they are critical for the association with the ORC complex and that their mutations impair and severely disrupt *Drosophila* development (BALASOV *et al*. 2009), (BLEICHERT *et al*. 2013), (BALASOV *et al*. 2015). In the current study, we found that M232 stabilizes the Orc6-ORC interaction in *Drosophila,* but this residue is absent in human protein. Notably, M232 is positioned at a distance favorable for sulfur-aromatic interaction with W228 (**Figure 3**), a type of interaction known to stabilize protein structures (VALLEY *et al*. 2012). Interestingly, introduction of Methionine at the corresponding position in human Orc6 (I239M) significantly improved the performance of the *Drosophila* ORC complex but had no positive effect on association with the human ORC complex in human cell lines. These findings suggest evolutionary diversification in the mechanisms of ORC assembly among Metazoan species.

Our previous work identified a highly positively charged motif RKRK (residues 198-202) localized within HsOrc6 C terminal domain just outside of HsOrc6 middle TFIIB-like domain, which is critical for HsOrc6/DNA binding (XU *et al*. 2020). Interestingly, this motif resembles a nuclear localization signal (NLS) (MAKKERH *et al*. 1996) and is conserved in humans and rodents but is not present in *Drosophila*. It appears that rather than increasing subunit affinity, mammals have evolved a strong nuclear localization signal (NLS) in Orc6 to increase its local concentration within the nucleus. Previous studies demonstrated that the Orc6 is transported into the nucleus where it is assembled into ORC complex (GHOSH *et al*. 2011). The presence of a strong NLS in mammals and amphibians, in which Orc6 displays weak association with the ORC complex, and its absence in *Drosophila*, where Orc6 binds the complex with high affinity, suggests that distinct molecular mechanisms have evolved to promote ORC assembly in different metazoan lineages. Our previous (XU *et al*. 2020) and presented here data suggest an interesting possibility that the presence of an NLS and an ability of HsOrc6 to bind DNA are necessary for the targeting of the “loose” human Orc6 to the nucleus and chromatin ultimately providing an additional anchoring for the ORC association with the DNA. We have shown before that during DNA interaction human Orc6 may induce a bend in DNA allowing for a tighter binding of the protein (XU *et al*. 2020). The functional six-subunit ORC complex in human cells is formed after core ORC1-5 joins the initial HsOrc6/DNA complex through the interaction of the α11 helix of HsOrc6 C-terminus (aa 231-242) with Orc3 (BLEICHERT *et al*. 2013) (BLEICHERT *et al*. 2015), (XU *et al*. 2020).

We propose that a strong NLS and enhanced Orc6-ORC complex affinity represents two independent strategies for facilitating ORC assembly. It is noteworthy that in our previous work we developed a human-*Drosophila* hybrid Orc6 containing both the human NLS and the *Drosophila* α11 helix. This hybrid protein fully substituted for the wild-type *Drosophila* Orc6 and was successfully used to model Meier-Gorlin syndrome in *Drosophila* (BALASOV *et al*. 2015), (BALASOV *et al*. 2020), (BALASOV *et al*. 2025). This functional compatibility of the human- *Drosophila* Orc6 chimera further supports the idea that alternative mechanisms can work together to achieve the same biological outcome - stable and functional ORC assembly required for DNA replication initiation.

## Materials and methods

### Mutagenesis

*Drosophila*- human hybrids were designed by using a PCR technique. All mutations were generated by site- direction mutagenesis following Stratagene’s (Agilent) protocol. ORC6 mutants and hybrids were cloned into the pUAS vector in frame with GFP under the *orc6* native promoter and injected into *w^1118^* fly embryos. The individual fly stocks were set up in background of *orc6^35^* deletion.

### Rescue of the Orc6 Mutants

For rescue experiments, 200- 300 progeny derived from heterozygous flies carrying the genotype *orc6^35^/Cy;GFP-Orc6^mut^* were scored for the presence of homozygous mutant adults *orc6^35^/orc6^35^;GFP-Orc6^mut^*. Rescue efficiency was calculated based on the expected segregation ratio and ranged from 10% to 30%.

### Immunoprecipitation (IP) from ovary

20 fresh dissected ovaries were crushed in the glass homogenizer with 100ul of high salt IP buffer (25mM Hepes 7.6, 12.5mM MgCl2, 100mM KCl, 0.1mM EDTA, 300mM NaCl, 0.01% Titon X100) and extracted for 1h at 4^0^ C with continuous rotation. Extract was centrifuged at 15,000g for 15 min. and supernatant diluted 3 times with regular IP buffer (25mM Hepes 7.6, 12.5mM MgCl2, 100mM KCl, 0.1mM EDTA, 0.01% Titon X100). Protein A-Sepharose (Bio Vision Cat.6501-5) and rabbit polyclonal ORC2 antibodies were added and incubated with supernatant overnight. Sepharose beads were washed 3 times with IP buffer (20-30 min.) and diluted in 10ul of IP buffer. Samples were boiled in loading buffer, separated in 10% SDS-polyacrylamide gel and transferred on Immobilon-P membrane (Millipore cat. IPVH00010).

### Immunoprecipitation and Immunoblotting from cell culture

Cells were lysed in IP buffer containing 50 mM HEPES (pH 7.5), 150 mM NaCl, 1 mM EDTA, 2.5 mM EGTA, 10% glycerol, 0.1% Tween-20, 1 mM DTT, and protease inhibitor cocktail (B14002, Selleck Chemicals). Lysates were sonicated and clarified by centrifugation. The resulting supernatants were incubated overnight at 4°C with GFP-Trap Magnetic Agarose (GTMA; Proteintech). After washing with IP buffer, bound proteins were eluted by boiling in SDS sample buffer and analyzed by SDS-PAGE and immunoblotting. Primary antibodies used in this study included an anti-ORC6 antibody described previously (MACHIDA *et al*. 2005b), anti-ORC2 antibody (sc- 32734; Santa Cruz Biotechnology), and anti-GFP antibody (sc-9996; Santa Cruz Biotechnology).

### Mitotic Chromosome Preparation

Preparation of mitoses was described previously (Lebedeva et al., 2000). Briefly, third instar larval neural ganglia were incubated in 0.075M KCl for 5 minutes, fixed in methanol with acetic acid (3:1) for 20 minutes and then dispersed in a drop of 50% propionic acid on a slide. Then slides were dried and stained with 5% Giemsa’s solution.

### Expression of nuclear localization signal mutant in salivary glands

The wild type human Orc6, *Drosophila* Orc6 or NLS mutant HsR198A/K199A/R200A/K201A were fused with GFP and cloned into pUAS vector under the control of the GAL4- inducible UAS promoter. Transgenic flies carrying these constructs were crossed with flies expressing GAL4 under the control of the Sgs3 promoter (Bloomington stock *w^1118^; P{w^+mC^= Sgs3-GAL4.PD}TP*). The Sgs3 promoter drives GAL4 expression specifically in the salivary glands of third-instar larvae. Salivary glands were dissected from third-instar larvae in PBS, and live fluorescence images were acquired within 40 minutes of dissection using a fluorescence microscope.

### Structure modeling and NLS prediction

Structural modeling of Orc3- Orc6 interaction was performed on COSMIC2 cloud platform for structural biology research using alhafold2 tool for multi- protein complexes (https://www.cosmic2.org/alphafold2).

For NLS prediction we used web based (https://nls-mapper.iab.keio.ac.jp) cNLS Mapper (KOSUGI *et al*. 2009b).

### Cell Culture and Viral Transduction

HCT116 p53−/− cells (BUNZ *et al*. 1998), generously provided by Fred Bunz (Johns Hopkins University), stably expressing dCas9-KRAB were maintained in McCoy’s 5A medium. An ORC6-targeting sgRNA (5ʹ- GTTGACCCGCGGCGTTCAC-3ʹ) was cloned into the pLRG lentiviral vector (a gift from the Vakoc laboratory, Cold Spring Harbor Laboratory), and lentivirus was produced using standard packaging procedures. GFP-tagged human ORC6 was cloned into an MSCV retroviral vector using In-Fusion cloning, and retrovirus was generated by co-transfection with packaging plasmids.

### MTT Assay

For growth assays, 5 × 10^5^ cells were plated in 6-well plates and transduced with ORC6 sgRNA lentivirus in the presence of 8 μg/mL polybrene. Two days after lentiviral transduction, cells were split at a 1:3 ratio and transduced with GFP-ORC6 retrovirus in the presence of 8 μg/mL polybrene. Puromycin selection was initiated 48 h after retroviral transduction. Four days after retroviral transduction, cells were harvested for immunoblot analysis, and 1,000 cells were seeded into 96-well plates for proliferation assays. Cell proliferation was measured every 24 h using the CellTiter 96® Non-Radioactive Cell Proliferation Assay (G4100, Promega) according to the manufacturer’s instructions. All experiments were performed in triplicate, and absorbance values were normalized to those obtained on the first day of measurement. Successful depletion of endogenous ORC6 and expression exogenous ORC6 was verified by western blot (**Supplementary figure 3**).

## Acknowledgments.

We would like to thank Jamie Roebuck, Model Systems Genomics, for Drosophila embryo injections.

---

**Supplementary Figure 1.**
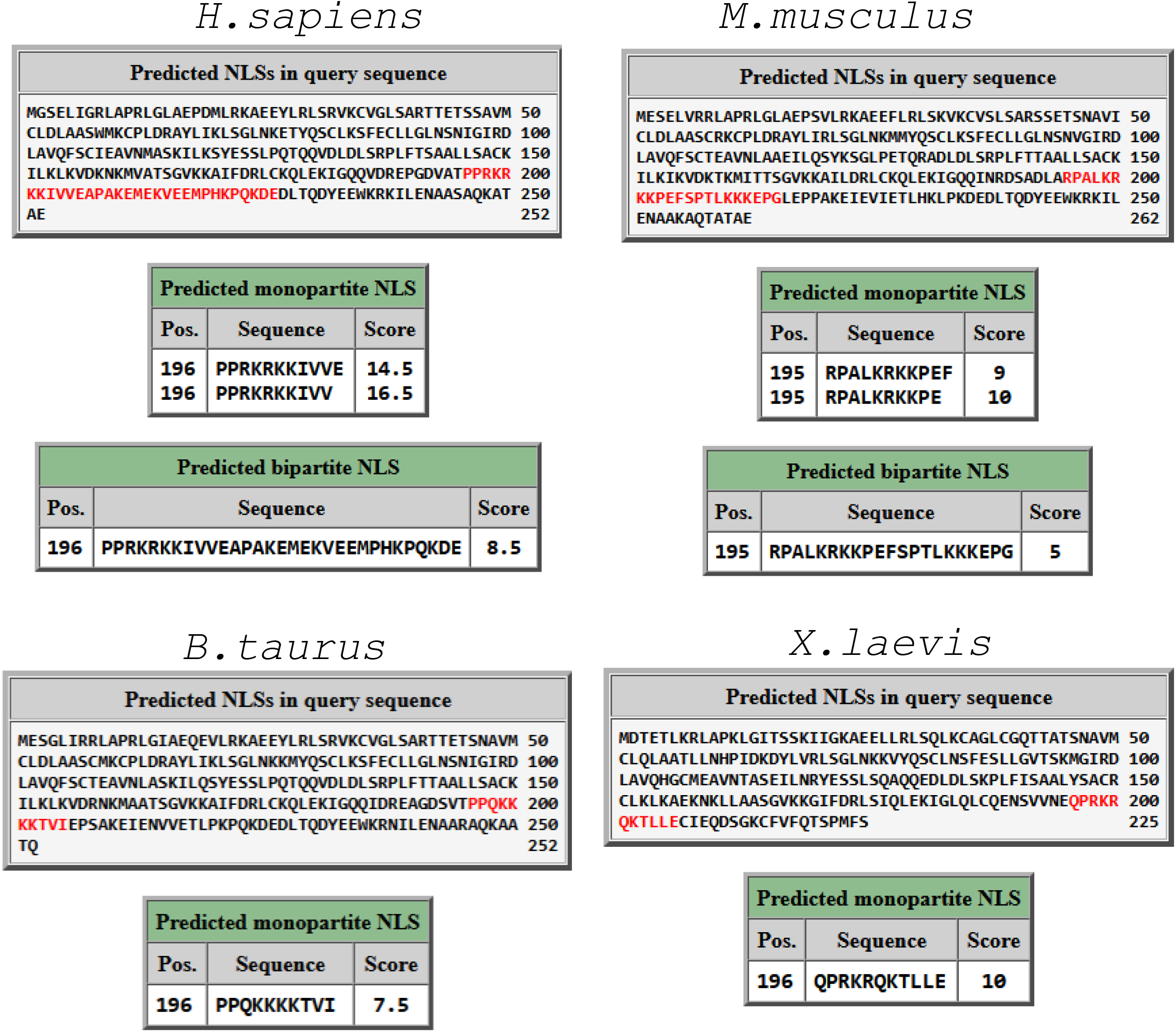
cNLS mapper results for human, bovine, mouse and xenopus Orc6. NLS with a score 7 and higher indicates exclusively nuclear localization and correlate with lose association Orc6 with Complex. All *Drosophila* species have score below 4 that indicate localization in both the nucleus and the cytoplasm. (https://nls-mapper.iab.keio.ac.jp)

**Supplementary Figure 2.**
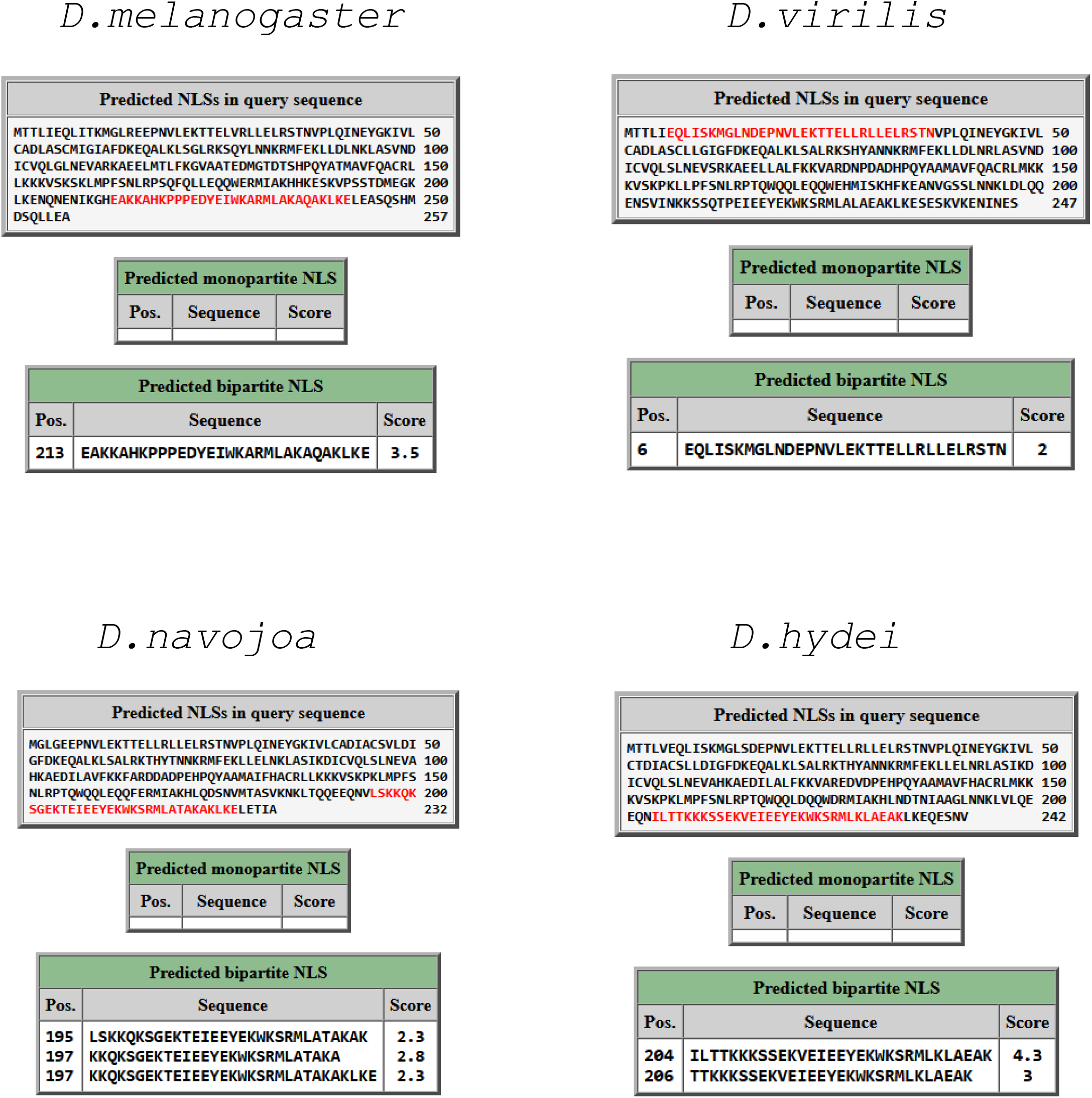
cNLS mapper results for *Drosophilidae*.

**Supplementary Figure 3.**
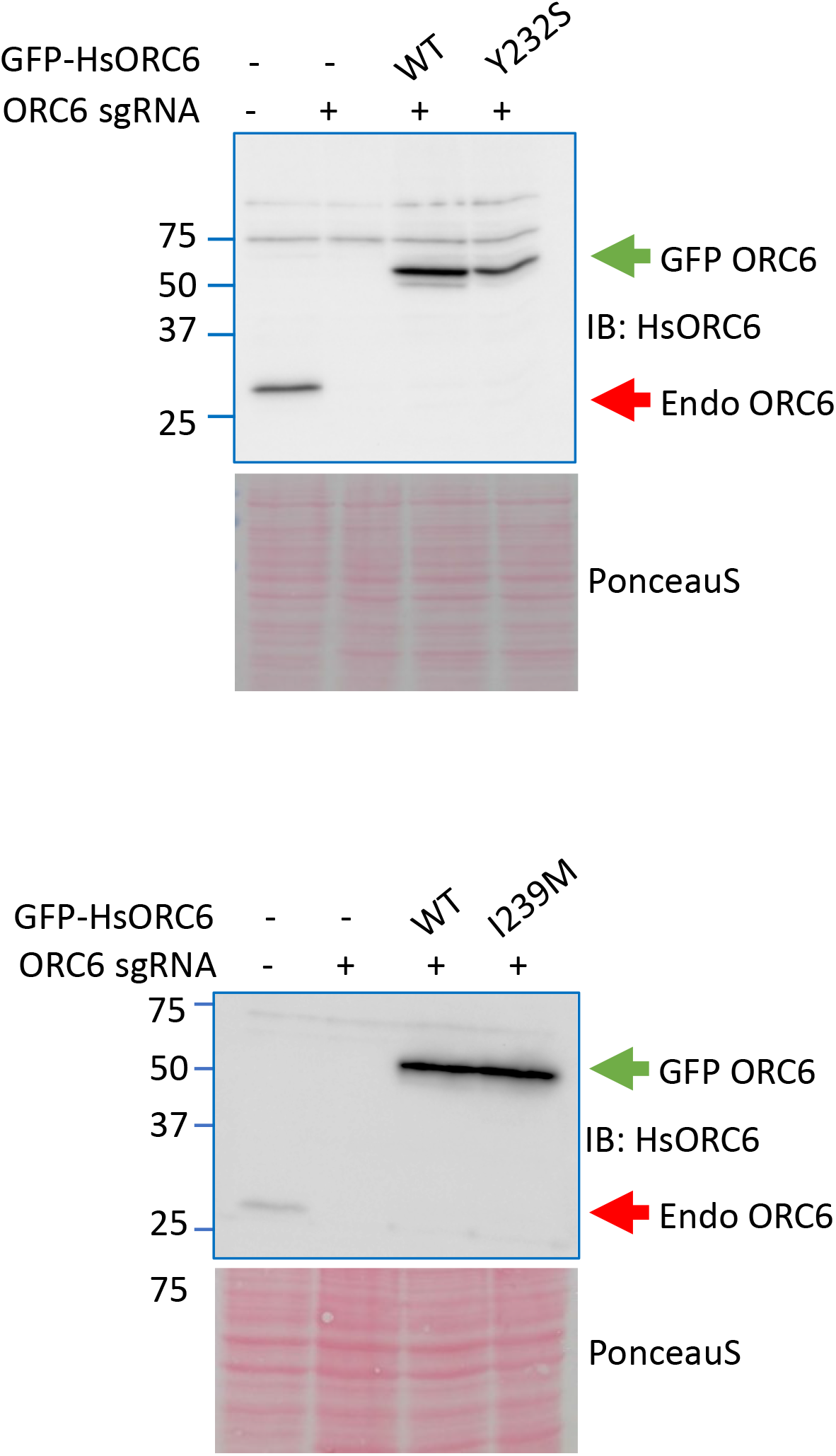
Western blot analysis of endogenous and exogenous ORC6 expression in cells depleted of endogenous ORC6 by CRISPRi and expressing GFP-tagged WT or mutant human ORC6 . Endogenous ORC6 and GFP-tagged ORC6 are indicated by red and green arrows, respectively. Ponceau S staining serves as a loading control.

